# Counting actin in contractile rings reveals novel contributions of cofilin and type II myosins to fission yeast cytokinesis

**DOI:** 10.1101/2021.07.26.453848

**Authors:** Mamata Malla, Thomas D. Pollard, Qian Chen

## Abstract

Cytokinesis by animals, fungi and amoebas depends on actomyosin contractile rings, which are stabilized by continuous turnover of actin filaments. Remarkably little is known about the amount of polymerized actin in contractile rings, so we used low concentration of GFP-Lifeact to count total polymerized actin molecules in the contractile rings of live fission yeast cells. Contractile rings of wild-type cells accumulated polymerized actin molecules at 4,900/min to a peak number of ∼198,000 followed by a loss of actin at 5,400/min throughout ring constriction. In *adf1-M3* mutant cells with cofilin that severs actin filaments poorly, contractile rings accumulated polymerized actin at twice the normal rate and eventually had almost two-fold more actin along with a proportional increase in type II myosins Myo2, Myp2 and formin Cdc12. Although 30% of *adf1-M3* mutant cells failed to constrict their rings fully, the rest lost actin from the rings at the wild-type rates. Mutations of type II myosins Myo2 and Myp2 reduced contractile ring actin filaments by half and slowed the rate of actin loss from the rings.

## Introduction

Cytokinesis separates daughter cells during the last stage of the cell cycle. Amoebas, fungi and animal cells assemble an actomyosin ring to provide force to form a cleavage furrow (for review see (Pollard and O’Shaughnessy, 2019)). Contractile rings consist of actin filaments, actin binding proteins including alpha-actinin, capping protein, cofilin, formins and type II myosins (Wu et al., 2003). In fission yeast, two type II myosins contribute about equally to the rate of ring constriction (Bezanilla et al., 2000; Laplante et al., 2015). Myo2 is essential for viability, while the unconventional myosin-II called Myp2 is not.

Although actin is the most abundant protein in contractile rings, much less is known about its dynamics than the myosins or actin-binding proteins owing to difficulty tracking actin in live cells. Fluorescent phalloidin is widely used to stain actin in fixed cells but this provides only a snapshot. SIR-actin, jasplakinolide conjugated to silicon rhodamine, can stain actin filaments in live cells (Lukinavicius et al., 2014), but jasplakinolide will alter actin dynamics. Microinjection of fluorescently labeled actin is an option for some animal cells (Cao and Wang, 1990) but has not been exploited for quantitative measurements and may not be feasible for some cells including fungi. Unfortunately, the genetically encoded fluorescent tags tested to date compromise the function of actin during cytokinesis. For example, the formins that nucleate and elongate actin filaments for the contractile ring in fission yeast filter out all the actin fused to either fluorescent proteins such as GFP or small tetracysteine peptide tags (Chen et al., 2012; Wu and Pollard, 2005).

Consequently, only indirect labeling of actin filaments in the contractile ring has been successful. Although less versatile than direct labeling, we measured about 190,000 actin molecules, equal to 500 µm of actin filaments, in the fission yeast contractile ring by titration with Lifeact (Courtemanche et al., 2016), a small peptide that binds actin filaments (Riedl et al., 2008). We also estimated the average length of the filaments from the ratio of polymerized actin to formin molecules in the contractile ring. The filaments start at about 1.4 µm and shorten gradually during the ring constriction. Despite this progress, little is known about how actin filaments themselves turn over in contractile rings.

Cofilins promote actin turnover during endocytosis in yeast (Chen and Pollard, 2013; Okreglak and Drubin, 2007), at the leading edge of motile cells (Ghosh et al., 2004; Konzok et al., 1999) and in neurite growth cones (Zhang et al., 2012). Cofilin is required for the assembly of the cytokinetic contractile ring in fission yeast (Chen and Pollard, 2011; Nakano and Mabuchi, 2006). Cofilin mutations that reduce severing activity slowed or prevented the assembly of contractile rings, because the precursors to the contractile ring, called cytokinetic nodes, are pulled into large heterogeneous clusters around the equator rather than being organized into a contractile ring. Rings that form slowly from these clusters constrict far more variably than those in wild-type cells (Chen and Pollard, 2011).

Here we expanded our previous study (Courtemanche et al., 2016) to count polymerized actin in contractile rings of wild-type fission yeast and strains with mutations in cofilin, formin and type II myosins. The cofilin mutant *adf1-M3* with half of wild-type severing activity had strong effects on both contractile ring assembly and disassembly. The contractile rings of the cofilin mutant cells have about twice the normal numbers of actin, formin Cdc12, Myo2 and Myp2 molecules, while contractile rings of cells with a hypomorphic mutation of formin Cdc12 accumulated actin slowly. Remarkably, mutations of either type II myosin, Myo2 or Myp2, reduced contractile ring actin by half.

## Results

### Measurement of actin turnover in contractile rings using GFP-Lifeact

To measure the number of polymerized actin molecules in contractile rings of live fission yeast over time, we expressed GFP-Lifeact (GFP-LA) constitutively from the endogenous *leu1* locus, driven by a constitutive *Padf1* promoter of the endogenous cofilin gene. All yeast strains used in this study expressed GFP-LA at similar levels (Fig. S1), which saturates only 6% of polymerized actin molecules and avoids artifacts during cytokinesis and endocytosis caused by higher concentrations of GFP-Lifeact (Courtemanche et al., 2016). Using a calibrated fluorescence microscope (Wu and Pollard, 2005), we converted the fluorescence intensity of GFP-Lifeact into the numbers of actin molecules in subcellular structures (Courtemanche et al., 2016). Specifically, we expressed from the endogenous locus Rlc1, the regulatory light chain for both Myo2 and Myp2, tagged with tdTomato and used its fluorescence to segment the contractile ring and measure the total GFP-Lifeact fluorescence.

Actin accumulated in contractile rings, about half during assembly and about half during 18 mins of maturation when the rate was constant at ∼4,900 molecules/min (Fig. 1A-B and Table 1). At the end of the maturation phase, just before cleavage furrow ingression, fully assembled contractile rings contained ∼200,000 polymerized actin molecules, consistent with previous measurements (Courtemanche et al., 2016).

**Figure 1.**
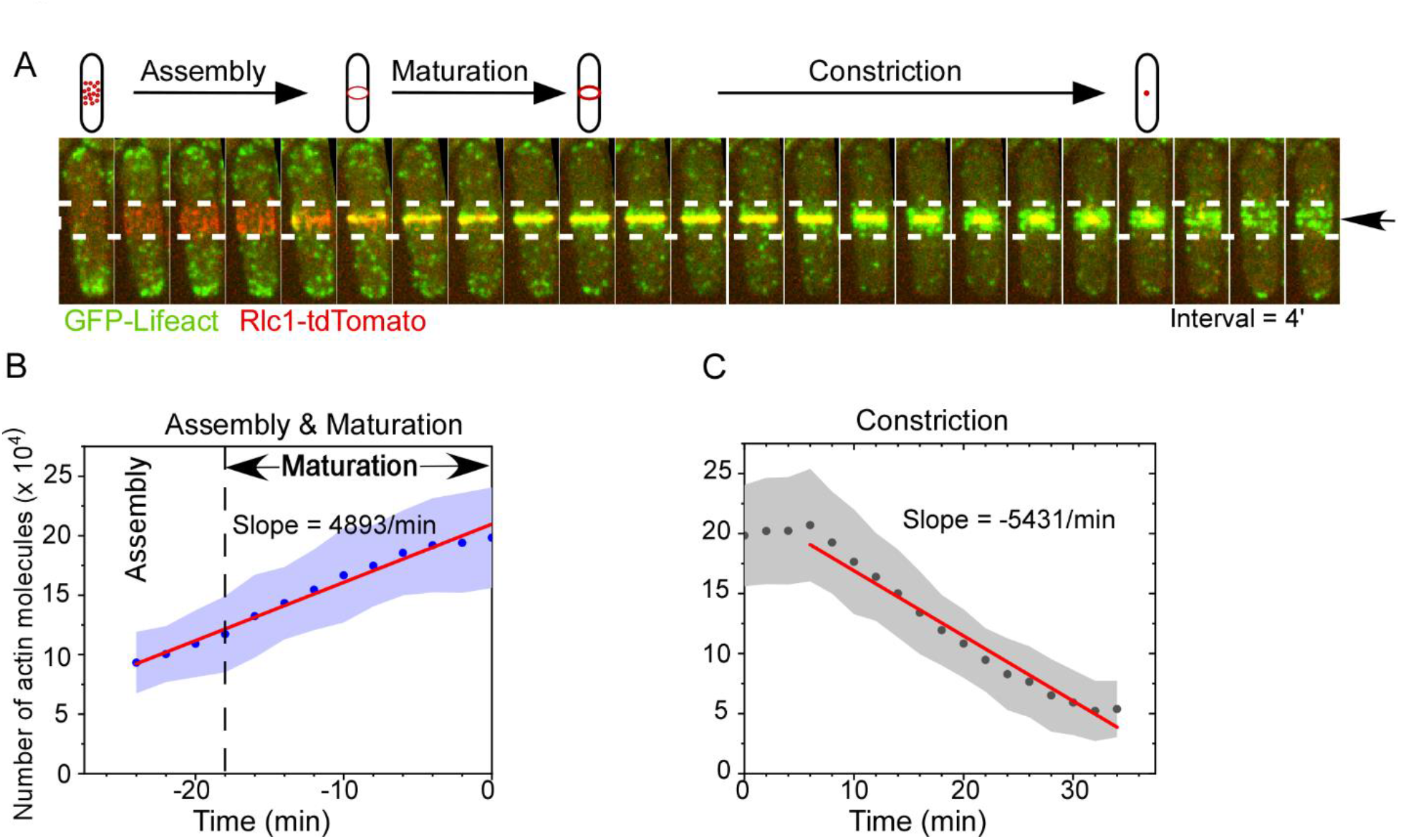
Counting the numbers of actin molecules in the contractile ring of fission yeast using GFP-Lifeact. **(A)** Assembly and constriction of the contractile ring documented by time-lapse micrographs of a cell expressing both GFP-Lifeact (green) and Rlc1-tdTomato (red). **(B-C)** Average time course of the number of actin molecules in the contractile rings during **(B)** their assembly and maturation and **(C)** constriction in wild-type cells. Clouds represent standard deviations. Time zero is defined as the start of the ring constriction. Red line represents the best linear fit. R^2^ > 0.9.

**Table 1:** Summary of the actin assembly and disassembly in the contractile ring.

| Genotype | Number of actin molecules in the mature ring ($\times 10^{-3}$)* | Number of cells measured* | Net assembly rate (actin molecules $\times 10^{-3}$ min<sup>-1</sup>)<sup>#</sup> | Net disassembly rate (actin molecules $\times 10^{-3}$ min<sup>-1</sup>)<sup>#</sup> | Normalized disassembly rate (min<sup>-1</sup>)<sup>&</sup> |
| --- | --- | --- | --- | --- | --- |
| *Wild-type* | 198 ± 42 | 80 | +4.9 | -5.4 | -2.7% |
| *adf1-M3* | 362 ± 176 | 21 | +10.0 | -5.8 | -1.6% |
| *cdc12-4A* | 93 ± 23 | 38 | +2.4 | N.M. | N.M. |
| *myo2-E1* | 82 ± 28 | 38 | N.M. | -1.6 | -2.0% |
| *myp2Δ* | 88 ± 21 | 33 | N.M. | -2.3 | -2.6% |

\*: The cells were pooled from at least two independent biological repeats.
<sup>#</sup>: Based on the best fits of linear regression. $R^2 > 0.90$ .
<sup>&</sup>: Equals to the net disassembly rate divided by the total number of actin molecules in the ring.
N.M.: Not measured.

During contractile ring constriction the number of polymerized actin molecules was constant for the first six minutes, which was not clear in our previous analysis (Courtemanche et al., 2016), and then declined steadily at ∼5,400 molecules/min for ∼20 minutes (Fig. 1C). About 50,000 actin molecules remained in the contractile ring remnant at the end of constriction, which has not been reported previously. These filaments abruptly dispersed within two minutes, together with the myosin regulatory light chain Rlc1 leaving behind actin patches along both sides of the cleavage furrow (Fig. 1A).

### Contractile ring assembly and composition in the cofilin hypomorphic mutant *adf1-M3*

Cofilin is essential for viability, so we used the hypomorphic mutant *adf1-M3* to test the role of cofilin in the dynamics of contractile ring actin filaments. The *adf1-M3* mutation reduces the severing activity of cofilin by more than 50% and slows contractile ring assembly (Chen and Pollard, 2011). Actin patches and actin filament bundles stained brighter with Bodipy-phallacidin in fixed *adf1-M3* mutant cells than in wild-type cells (Chen and Pollard, 2011). However, at that time we lacked quantitative probes to measure polymerized actin in live cells (Chen et al., 2012). The cytokinesis defects were less severe in the *adf1-M3* strain than the *adf1-M2* strain (Chen and Pollard, 2011), allowing us to analyze the assembly and disassembly of larger numbers of mature rings.

Contractile ring assembly in *adf1-M3* mutant cells was less orderly (Fig. 2B) and much more variable than in wild type cells (Fig. 2B-D). Contractile rings of *adf1-M3* mutant cells accumulated actin twice as fast over a similar period of time as wild type cells (Table 2). Therefore, mature rings of the mutant had on average about 1.9 times as much actin as wild-type cells (Table 1 and Fig. 2A, arrows). The peak number of molecules (100,000 to 600,000) was much more variable in *adf1-M3* mutant cells than wild-type cells (Fig. 2D). On average, the contractile rings of *adf1-M3* mutant cells had enough actin molecules to assemble ∼ 950 µm of filaments (Table 2). A cross section of such rings would contain ∼75 filaments in a bundle ∼160 nm wide, if the spacing between the actin filaments is 15 nm like wild-type cells, which have ∼50 filaments in a bundle of ∼125 nm wide (Courtemanche et al., 2016; Swulius et al., 2018). We conclude that severing by cofilin limits and makes more reliable the assembly of actin filaments in the contractile ring.

**Figure 2.**
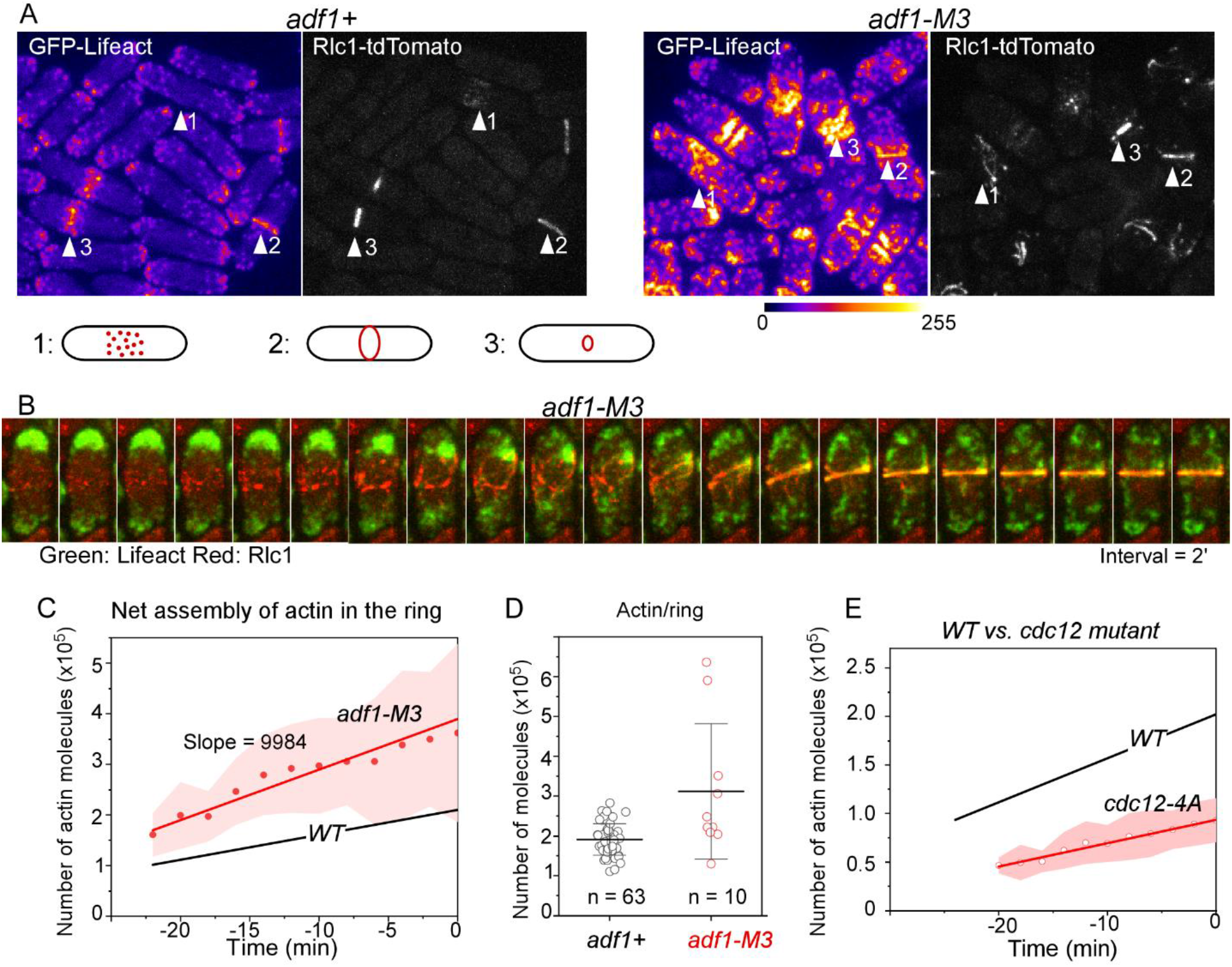
Contractile ring assembly in cofilin mutant *adf1-M3* and formin mutant *cdc12-4A*. **(A)** Fluorescence micrographs of the wild-type (left) and *adf1-M3* cells. Both expressed GFP-Lifeact (fire colored) and Rlc1 -tdTomato. Arrowheads: contractile rings at the stages of (1) assembly, (2) maturation and (3) constriction. **(B)** Time series of micrographs (pseudo-colored) of a cofilin mutant cell during cytokinesis. **(C)** Time course of actin molecules accumulating in the contractile rings of *adf1-M3* cells (n = 21). Red symbols are mean values; the line represents best fit; the cloud represents standard deviations. The line for *WT* cells is from Fig. 1B. **(D)** Average number of actin molecules in mature contractile rings. Measurements were taken just before the start of the ring constriction. **(E)** Time course of actin assembly in the contractile rings of formin *cdc12-4A* mutants. The line for *WT* cells is from Fig. 1B. The slope of the *cdc12-4A* mutant cells is significantly smaller than that of the wild-type (P < 0.005).

**Table 2:** Comparison of the architecture of the contractile ring between wild-type and the cofilin mutant.

| Genotype | Estimated assembly time of actin (mins)* | Total filament length in the mature ring ($\mu\text{m}$) | Number of formin dimers in the mature ring | Number of type II myosin motor domains in the mature ring | Average filament length in the mature ring ($\mu\text{m}$) |
| --- | --- | --- | --- | --- | --- |
| *WT* | 43 | 530 | ~350 | ~7,000 | 1.5 |
| *adf1-M3* | 39 | 975 | ~520 | ~14,000 | 1.9 |

\*: Based on the best fit of linear regression. Time = Number of actin molecules/net assembly rate.

To estimate the lengths of actin filaments in the contractile ring of *adf1-M3* mutant cells we measured the number of formins in fully assembled contractile rings just before they constricted. Assuming that barbed ends of all contractile ring actin filaments are associated with a formin (Coffman et al., 2013), the ratio of total actin to total formin molecules gives the average length of actin filaments, averaging ∼1.5 µm in wild-type cells (Table 2), similar to the previous estimate (Courtemanche et al., 2016).

The number of Cdc12-3GFP molecules in the contractile rings of *adf1-M3* mutant cells was on average about twice that of wild-type cells and much more variable (Fig. 3C), while the number of formin For3 was the same as wild-type cells (Fig. 3D). As a result, the combined number of the two formin molecules was ∼50% higher in the contractile rings of the mutant cells (Table 2). Consequently, the ratio of actin to formins was ∼25% higher, translating to an average length of 1.9 µm in the mutant cells (Table 2).

**Figure 3.**
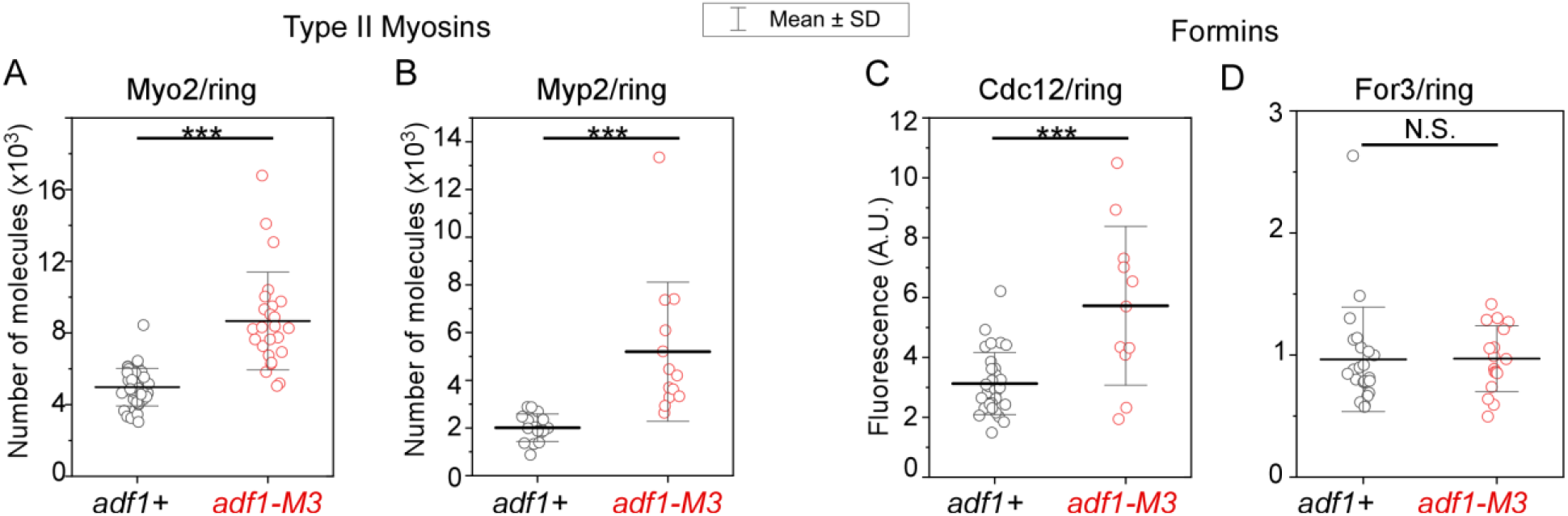
Architecture of the contractile ring in cofilin mutant cells. **(A-B)** Numbers of type II myosins, **(A)** Myo2 and **(B)** Myp2 in contractile rings of wild-type and *adf1-M3* cells. **(C-D)** The fluorescence intensities of two formins, **(C)** Cdc12-3GFP and **(D)** For3-3GFP in contractile rings of wild-type and *adf1-M3* cells. ***: P < 0.001. N.S.: P > 0.05. Values were calculated from two-tailed student t-tests.

The numbers of type II myosins (Myo2 and Myp2) in mature contractile rings of wild-type cells peaked at ∼7,000 myosin molecules (Fig. 3A-B and Table 2), consistent with previous measurements (Goss et al., 2014; Wu and Pollard, 2005). This translates to one myosin motor domain for every 76 nm of actin filament and ∼20 motor domains for every actin filament in the contractile ring.

Contractile rings of *adf1-M3* mutant cells that were able to constrict had twice as many myosin molecules as the wild-type cells, translating to one myosin motor domain for every 70 nm of filament (Fig. 3A-B and Table 2). We conclude that reduced severing by cofilin leads to contractile rings with twice the normal numbers of actin, Cdc12 and type II myosin molecules, so the ratios of these three proteins are about the same as in wild-type cells (Table 2).

### Effects of a formin mutation on contractile ring assembly

Measurements of the time course of actin accumulation in the contractile rings of the *cdc12-4A* formin mutant cells revealed defects in actin that were not appreciated in previous work (Bohnert et al., 2013). The essential formin Cdc12 is required for assembly of actin filaments in contractile rings (Chang et al., 1996; Kovar et al., 2003). The hypomorphic *cdc12-4A* mutation prevents the phosphorylation of the formin by the essential SIN pathway kinase Sid2 (Bohnert et al., 2013). In prior work the mutant cells appeared to assemble normal contractile rings, but our quantitative measurements revealed that the *cdc12-4A* mutation reduced by about half both the rate of accumulation and the final numbers of polymerized actin (Fig. 2E and Table 1).

### Contractile ring disassembly during constriction in *adf1-M3* mutant cells

The contractile rings in 30% of *adf1-M3* mutant cells (n = 65) either failed to constrict or halted constriction prematurely (Fig. S2), but those that constricted fully did so at an average rate similar to wild-type cells although with much more variability (Chen and Pollard, 2011). During ring constriction, the absolute number of actin molecules in the contractile rings of *adf1-M3* cells declined linearly at ∼5,800/min (Fig. 4A and B, Table 1), almost identical to wild-type cells. However, the cofilin mutant cells had two defects. First, the large standard deviations showed that rate of actin disassembly varied much more in *adf1-M3* cells than in wild-type cells. Second, the normalized disassembly rate, which took the number of actin molecules in the ring into consideration, was 40% lower in the mutant than wild-type cells. Thus, normal severing by cofilin is not essential for the disassembly of actin filaments in constricting contractile rings but makes the process much more orderly.

**Figure 4.**
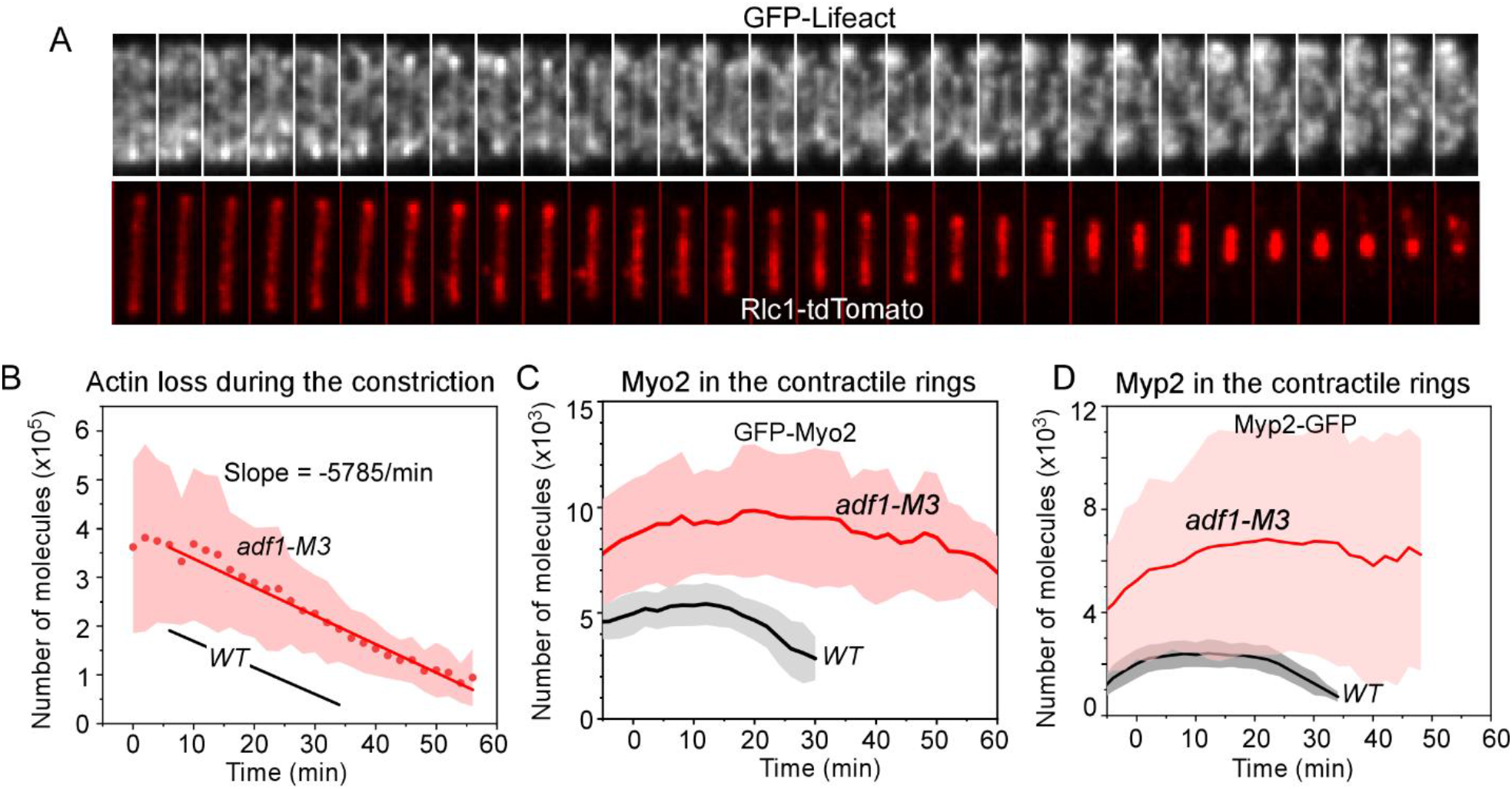
Time courses of the numbers of actin and type II myosins in constricting contractile rings of WT and *adf1-M3* cells. Time zero is the onset of constriction. **(A)** Time-series of micrographs of a constricting contractile ring in an *adf1-M3* mutant cell expressing GFP-Lifeact (top) and Rlc1-tdTomato (bottom, red). Interval = 2 min. **(B)** Time course of the loss of actin molecules from contractile rings as they constricted. Red symbols and line are mean values; the cloud shows standard deviations. **(C)** Time course of the numbers of GFP-Myo2 molecules in constricting contractile rings of WT and *adf1-M3* cells (n> 25). Red symbols and line are mean values; the cloud shows standard deviations, which are large in the mutant cells. **(D)** Time course of the numbers of Myp2-GFP molecules in constricting contractile rings of WT and *adf1-M3* cells (n > 10). The thick lines are mean values.

Wild-type cells retain both Myo2 (Fig. 4C) and Myp2 (Fig. 4D) in the contractile ring through the first 20 minutes of constriction before they leave during the last 10 minutes of the constriction as observed earlier (Wu and Pollard, 2005). In contrast, these myosins persisted at nearly their highest levels for an hour and the time course of the process was much more variable in the *adf1-M3* mutant cells (Fig. 4C and D). In a few cofilin mutant cells, the myosins dwelled at the cell division for more than 10 minutes after the completion of the ring constriction (Fig. 5A). Myp2 oscillated as clusters along the contractile ring of the mutant cells, which was rarely observed in the wild-type cells (Fig. 5A).

**Figure 5.**
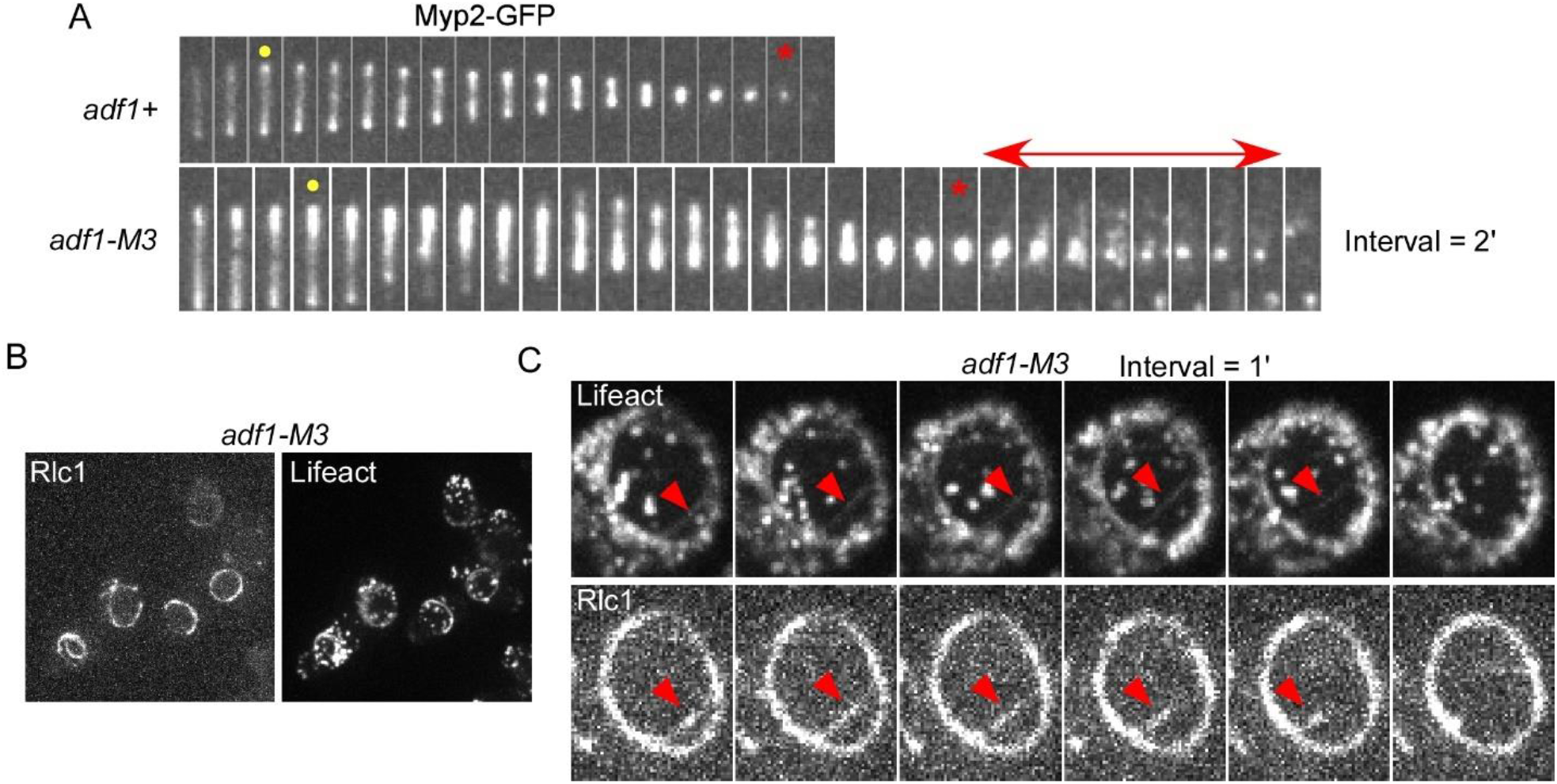
Disassembly of myosins in the contractile ring of the cofilin mutant. **(A)** Time series of micrographs of contractile rings in a wild-type and an *adf1-M3* cells expressing Myp2-GFP. Dot: Start of ring constriction. Asterisk: Completion of ring constriction. **(B)** Micrographs of attached *adf1-M3* cells in a petri dish. The spots around the contractile ring are actin patches. **(C)** Time series of micrographs of the contractile ring in a cofilin mutant cell. Arrowhead: shedding actin filaments (upper panel) and myosin (lower panel).

Face views of constricting contractile rings of *adf1-M3* cells revealed linear structures containing both myosin-II isoforms and actin filaments that separated from 60% (n = 18) of the rings (Fig. 5B and C). Similar structures were observed previously in 3D-reconstructions of cofilin mutants *adf1-M2* and *-M3* (Cheffings et al., 2019; Chen and Pollard, 2011). Although the structures containing myosin-II retracted back to the contractile ring, the bundles of actin filaments did not (Fig. 5C, arrowheads). Such shedding of actin filaments was not observed in the wild-type cells. We conclude that contractile rings are far less stable in cofilin mutant cells than wild type cells.

### Influence of type II myosins on assembly and disassembly of contractile ring actin filaments

Mutations of either type II myosin gene in the *myo2-E1* or *myp2Δ* strains reduced the numbers of actin molecules in contractile rings by more than half compared with wild-type cells at the end of the maturation period and the onset of constriction (Fig. 6A-B and Table 1). This surprising finding was missed previously.

**Figure 6.**
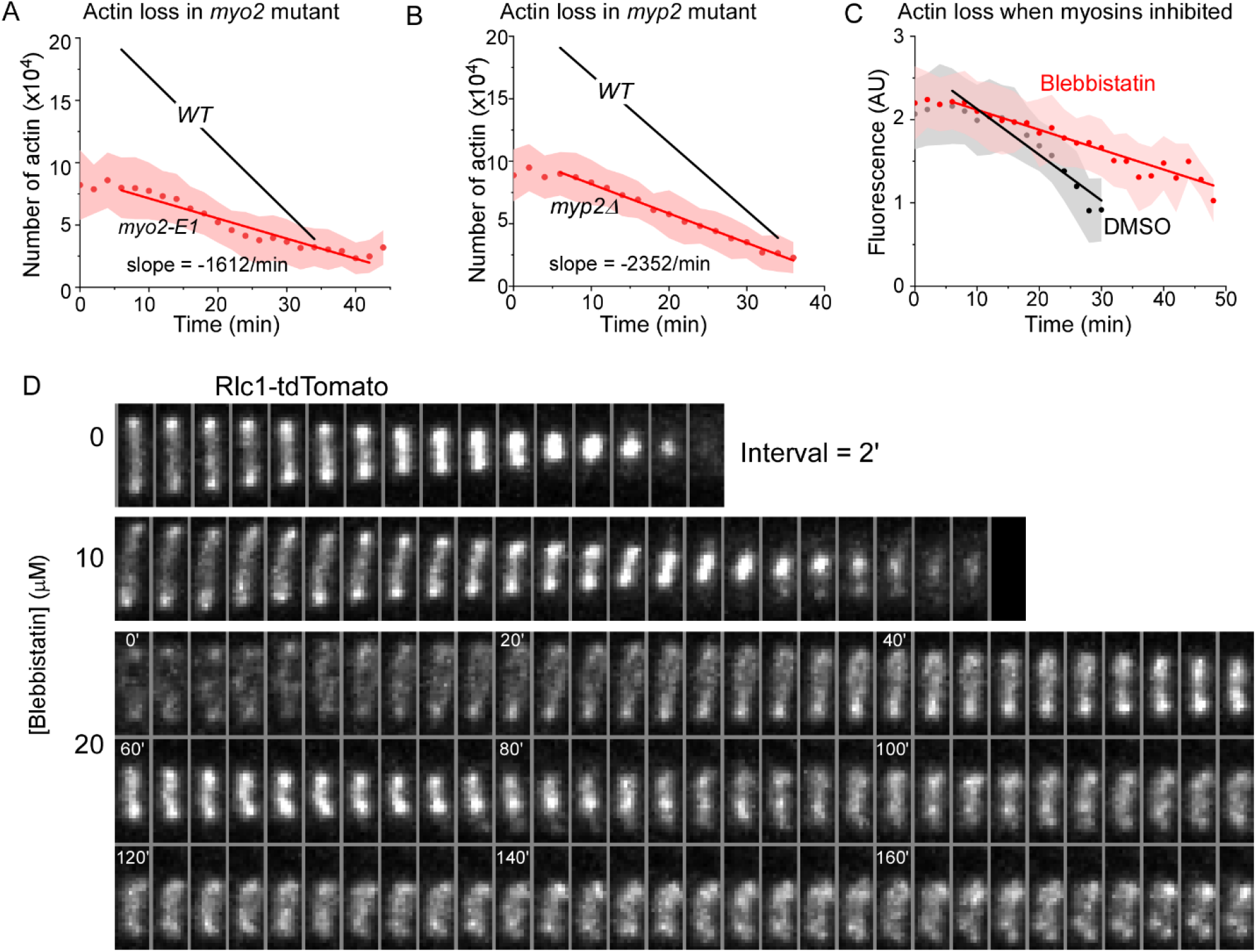
Disassembly of contractile ring actin filaments in type II myosin mutants at the permissive temperature of 22°C. **(A-B)** Time courses of actin loss from constricting contractile rings of **(A)** *myo2-E1* and **(B)** *myp2Δ* mutant cells. Dark red circles are mean values; red lines are the best linear fits; the cloud shows standard deviations. **(C)** Time courses of GFP-Lifeact fluorescence in contractile rings of the cells treated with either 10 µM blebbistatin or 1% DMSO (control). The slopes are -5489/min (DMSO) and-2389/min (Blebbistatin) respectively. Only cells with a mature contractile ring before the treatment were included. Circles are mean values; solid lines are the best linear fits; the cloud shows standard deviations. **(D)** Time series of micrographs of contractile rings marked by Rlc1-tdTomato in three wild-type cells treated with either DMSO (top), or 10 (middle) or 20 µM (bottom) of blebbistatin. The number in the time series of 20 µM blebbistatin-treated cell represents time in minutes.

Starting with less than the normal amount of polymerized actin, contractile rings constricted slower in both *myo2-E1* (0.32 µm/min) and *myp2Δ* (0.30 µm/min) cells than wild-type cells (0.36 µm/min). These rates are slightly higher than earlier studies (Laplante et al., 2015; Zambon et al., 2017). After normalization for the initial actin content, the rates that actin left constricting rings were only slightly less than normal in the *myo2-E1* mutant and similar in the *myp2Δ* strain to wild-type cells (Table 2).

To rule out the possibility that the lower disassembly rates in the myosin mutants were indirectly tied to ring assembly defects, we measured the loss of actin from contractile rings of wild-type cells treated with blebbistatin to inhibit myosin II (Straight et al., 2003). Contractile rings in wild-type cells either disintegrated or failed to assemble in blebbistatin concentrations of ≥20 µM (Fig. 6D). Treating wild-type cells with mature contractile rings with 10 µM blebbistatin decreased the rate of actin disassembly by 60% (Fig. 6C). The ring constriction rate was also lower by 30% (n = 16) (Fig. 6D). We conclude that type II myosins contribute to both the assembly and disassembly of actin filaments in contractile rings.

## Discussion

Knowing the numbers and dynamics of polymerized actin molecules in the actomyosin contractile ring is essential for understanding the mechanism of cytokinesis, but such measurements have not been made due to the technical challenge of labeling actin directly without disrupting its activity (Chen et al., 2012; Wu and Pollard, 2005). Indirect probes can produce artifacts, so we measured polymerized actin with a low concentration of GFP-Lifeact that does not disturb endocytosis or cytokinesis (Courtemanche et al., 2016). Measurements on beautiful EM-tomograms (Swulius et al., 2018) confirmed our earlier count of contractile ring actin with GFP-Lifeact (Courtemanche et al., 2016).

Our method provides valuable, new quantitative data on the accumulation and loss of polymerized actin in contractile rings but does not reveal the behavior, including the turnover, of individual filaments. Experiments with a probe directly on actin molecules will be required to probe the underlying mechanisms.

Our measurements revealed several aspects of cytokinesis that were overlooked due to the lack of quantitative data on actin in contractile rings including the phenotypes of yeast strains with mutations in cofilin, myosin and formin genes. The protein products of each of these genes are essential for cytokinesis and their roles are likely to have been conserved during evolution.

### The role of cofilin in the assembly and composition of contractile rings

A mutation that reduces the severing activity of cofilin has a remarkable impact on the molecular composition of the contractile ring: about twice the wild-type numbers of polymerized actin, Myo2, Myp2 and formin Cdc12. First we consider myosins and then formins and actin for discussion.

#### Myosins

Myo2 and formin Cdc12 are components of cytokinesis nodes that form prior to the assembly of actin filaments (Wu and Pollard, 2005), so extra numbers of both proteins in the contractile rings of *adf1-M3* cells implies proportionally more cytokinesis nodes. In fact, the nodes were larger in *adf1-M3* mutant cells than wild-type cells (Chen and Pollard, 2011) and likely represent clusters of larger numbers of small, unitary nodes as revealed in wild-type cells by super-resolution microscopy (Laplante et al., 2016). We do not know when these extra nodes form or how either reduced severing or more polymerized actin induce their formation.

On the other hand, Myp2 is recruited to fully formed contractile rings during the maturation period in a process that depends on actin filaments (Okada et al., 2019; Takaine et al., 2015; Wu et al., 2003). Thus, the higher numbers of Myp2 molecules in contractile ring of *adf1-M3* cells may follow directly from the high content of actin filaments.

#### Formins and actin

The essential formin Cdc12 nucleates and elongates actin filaments in the contractile ring (Chang et al., 1996; Kovar et al., 2003), so more formin Cdc12 in contractile rings of *adf1-M3* cells likely contributes to the rapid accumulation of extra actin filaments and the actin monomer concentration in the cytoplasm sets the rate of growth of individual filaments. On the other hand, no connection between slow severing and excess formin is known.

The longer actin filaments in the contractile rings of *adf1-M3* cells are expected for cells with low actin filament severing activity (Chen and Pollard, 2011) and given evidence that cofilin stochastically severs actin filaments connecting the precursor nodes (Chen and Pollard, 2011). Longer filaments may also explain our observation of a negative genetic interaction between cofilin and *ain1* mutants (Chen and Pollard, 2011).

Since actin filaments accumulate faster in the contractile rings of *adf1-M3* mutant cells, some other mechanism must account for the slow assembly of full contractile rings (Chen and Pollard, 2011). The most likely mechanism is that slow severing results in the aggregation of nodes that delays the coalescence of the nodes into an organized contractile ring of actin oligomers (Chen and Pollard, 2011).

In contrast to the essential formin Cdc12, *adf1-M3* cells do not accumulate excess formin For3, which is not a component of cytokinesis nodes and joins fully assembled contractile rings at a later stage and assembles peripheral actin bundles at the cell division plane (Coffman et al., 2013). Unlike Cdc12, its recruitment likely depends on the type V myosin and polarized membrane secretion (Coffman et al., 2013).

### The role of cofilin in contractile ring constriction and disassembly

Multiple defects appear during constriction of contractile rings of *adf1-M3* cells. First, constriction is much less uniform than in wild-type cells. About 30% of the rings fail to complete the constriction. Bundles of actin filaments containing myosins peel from the rings of mutant cells. Second, starting with contractile rings containing much more polymerized actin and both type II myosins, the cofilin mutant cells required more time for all three proteins to leave the rings in a highly variable fashion. Third, type II myosin Myp2 oscillates along the contractile rings as clusters. Nevertheless, the *adf1-M3* mutation does not significantly reduce the rate of net loss of actin filaments from contractile rings as they constrict.

The most likely explanation for the variability of contractile ring constriction in the *adf1-M3* cells is that the loss of severing activity compromises the continuous, relatively rapid (tens of seconds) turnover of contractile ring components, which is required to maintain orderly force production in computer simulations of constriction (Stachowiak et al., 2014). Those simulations assume estimates of continuous protein turnover revealed by photobleaching experiments. Dialing down the turnover of polymerized actin, Cdc12 and Myo2 in these simulations, resulted in the loss of tension in less than 3 minutes. Cofilin contributes to this turnover by severing actin filaments, while other uncharacterized processes cause the exchange of Cdc12 and Myo2 with cytoplasmic pools.

### Role of formins in the assembly of the contractile ring

Counting polymerized actin revealed unrecognized defects caused by the *cdc12-4A* mutation. This mutation prevents phosphorylation of Cdc12 by the SIN pathway kinase Sid2, but does not cause any discernable cytokinetic defects (Bohnert et al., 2013). Although cytokinesis appears normal, contractile rings in *cdc12-4A* mutant cells accumulate actin slower and in less than half the number of wild-type cells. This previously overlooked actin assembly defect may explain the hyper-sensitivity of this strain to Latrunculin A (Bohnert et al., 2013) as well as the genetic interaction of the *cdc12-4A* mutation with many other cytokinetic mutants.

### Roles of type II myosins in the assembly of the contractile ring

Our experiments revealed to our surprise that mutations of type II myosins strongly reduce the actin content of contractile rings. This remarkable, unexpected finding had been missed in dozens of studies on these mutant strains due to the lack of methods to measure actin in live cells. During ring assembly, Myo2 in a given node pulls on actin filaments growing from nearby nodes (Vavylonis et al., 2008). In a reconstituted system, force on a filament growing from formin Cdc12 slows its elongation (Zimmermann et al., 2017). However, the loss of Myo2 activity in the *myo2-E1* mutant has the opposite effect, resulting in more actin filaments, so the mechanism reducing the actin content is not clear. Myp2 is not present in contractile rings until after they form, so it must have its effect on the half of the actin that assembles during the maturation period, which is slower in the *myp2Δ* mutant than wild-type cells.

### The role of type II myosins in constriction and disassembly of the contractile ring

Contractile rings with half the normal amount of actin filaments constrict 10% slower in the *myo2-E1* strain and 20% slower in the *myp2Δ* strain than wild-type cells (Laplante et al., 2015). Disassembly of actin filaments is slower in both myosin mutants than wild-type cells, although these rates are the same as wild-type when normalized for the starting actin content. Previously, the defects in ring constriction in the strains with myosin-II mutations was attributed entirely to loss of function of the myosins (Laplante et al., 2015; Zambon et al., 2017). However, these strains have a secondary defect, a loss of about half the normal polymerized actin. This insight emphasizes the importance of quantitative measurements of other contractile ring components in mutant strains.

Inhibition of myosins with 10 µM blebbistatin reduced both the disassembly rate of actin filaments and the ring constriction rate by ∼60% and 30% respectively. In addition to cofilin-mediated severing, forces produced by myosins may contribute to actin filament turnover during contractile ring constriction.

Collectively, our experiments demonstrate the value of measurements of polymerized actin in live cells. These measurements provided new, unanticipated features of contractile ring assembly and constriction, which will motivate future studies to characterize mechanisms.

## Materials and methods

### Yeast genetics

We followed the standard protocols for yeast cell culture and genetics. Tetrads were dissected using a SporePlay+ dissection microscope (Singer, UK). Table 3 lists all the strains used in this study.

**Table 3.** Yeast strains.

| Strains | Genotype | Source |
| --- | --- | --- |
| QC-Y601 | *h+ leu2::kanMX-Pcof1-mEGFP-Lifeact rlc1-tdTomato-Nat* | Lab stock |
| QC-Y620 | *h+ leu2::kanMX-Pcof1-mEGFP-Lifeact rlc1-tdTomato-Nat adf1::KanMX6-Padf1-*adf1*-M3* | Lab stock |
| QC-Y1164 | *h? leu2::kanMX-Pcof1-mEGFP-Lifeact rlc1-tdTomato-Nat Myo2-E1* | This study |
| QC-Y1185 | *h? leu2::kanMX-Pcof1-mEGFP-Lifeact rlc1-tdTomato-Nat cdc12-4A::kanR ade6-M21X leu1-32 ura4-D18* | This study |
| QC-Y1196 | *h? leu2::kanMX-Pcof1-mEGFP-Lifeact rlc1-tdTomato-Nat myp2::kanMX6* | This study |
| JW766 | *h+ kanMX6-Pmyo2-GFP-myo2 ade6-M210 leu1-32 ura4-D18* | Lab stock<br>(Jian-Qiu Wu) |
| QC-Y129 | *h+ kanMX6-Pmyo2-GFP-myo2 leu1-32 ura4-D18 adf1M3A-KanMX6* | Lab stock |
| KV344 | *h? cdc12-3XGFP::kanMX6 leu1-32 his3-D1 ura4-D18 ade6-M216* | Lab stock<br>(David Kovar) |
| QC-Y142 | *h+ cdc12-3XGFP::kanMX6 leu1-32 his3-D1 ura4-D18 ade6-M216 adf1M3-KanMX6 leu1-32 ura4-D18 ade6-M216* | Lab stock |
| QC-Y267 | *h+ for3-3GFP-ura4+ ade6-M21 leu1-32 ura4-D18* | Lab stock |
| QC-Y1355 | *h? adf1M3A-KanMX6 leu1-32 ura4-D18 his3-D1 ade6-M216 for3-3GFP-ura4+ ade6-M21 leu1-32 ura4-D18* | This study |
| QC-Y519 | *h- myp2-GFP-kanMX6 ade6-M210 leu1-32 ura4-D18* | Lab stock |
| QC-Y1371 | *h? adf1M3A-KanMX6 leu1-32 ura4-D18 his3-D1 ade6-M216 myp2-mEGFP-KanMX6 ade6-M210 leu1-32 ura4-D18* | This study |

### Microscopy

For microscopy, 1 ml of exponentially growing yeast cells at 25°C with a density between 5.0 × 10^6^/ml and 1.0 × 10^7^/ml in YE5s liquid media (unless specified), were harvested by centrifugation at 4000 rpm for 1 min and resuspended in 50 µl YE5s. Due to the extremely slow growth of the *adf1-M*3 cells at 25°C, they were first inoculated at 30°C for a day from the plate, before being inoculated at 25°C for 12 h. 6 μL of re-suspended cells were applied to a 25% gelatin + YE5s pad and sealed under the coverslip with VALAP (a mix of an equal amounts of Vaseline, Lanolin, and paraffin) (Wang et al., 2016). Live cell microscopy was carried out on an Olympus IX71 microscope equipped with both a 100× (NA = 1.41) and a 60× (NA = 1.40) objective lenses, a confocal spinning-disk unit (CSU-X1; Yokogawa, Japan), a motorized XY stage with a Piezo Z Top plate (ASI). The images were captured on an Ixon-897 EMCCD camera controlled by iQ3.0 (Andor, Ireland). Solid-state lasers of 488 nm (5% for 100ms except for For3-3GFP: 7.5% for 150 ms) and 561 nm (5% for 50 ms) were used in the confocal fluorescence microscope. In all experiments, the cells were imaged for 3 h (2-minutes interval unless specified) by acquiring a Z-series of 15 slices at a step size of 0.5 μm using the 100× objective (unless specified). Live cell microscopy was conducted in a room where the temperature was maintained at around 22 ± 2°C. To minimize the variations in culture and microscopy conditions, we imaged both control and experimental groups with randomized order within a week. To measure the relative fluorescence of Cdc12-3GFP and For3-3GFP, a Z-series of 8 slices with a spacing of 1 μm was used.

To image the contractile ring head-on, cofilin mutant cells were observed in a petri dish with a coverslip bottom. A sample of 20 µl of the cell culture was spotted onto a glass coverslip (#1.5) at the bottom of a 10-mm petri dish (Cellvis, USA). The coverslip was precoated with 50 µl of 50 µg/ml lectin (Sigma, L2380) and allowed to dry overnight at 4°C. The cells were allowed to attach to the lectin for 10 minutes at room temperature. YE5s medium (2 ml) was then added to the dish just before microscopy. Images were acquired at 1-minute intervals.

### Drug treatment

A sample of 20 µl of the cell culture was spotted onto a glass coverslip on the bottom of a petri dish as described above. YE5s medium (2 ml) with either 10 µM Blebbistatin or 1% DMSO was added to the dish before starting microscopy using the 60x objective. Image acquisition was started exactly after 15 min of the drug treatment to minimize experimental variability.

### Image processing and analysis

We used ImageJ (National Institutes of Health) to process all the images, with either freely available or customized macros/plug-ins. For quantitative analysis, the fluorescence micrographs were corrected for X-Y drifting using the StackReg plug-in (Thevenaz et al., 1998). Average intensity projections of Z-slices were used for actin number measurements. Maximum intensity projections of Z-slices were used for the contractile ring tracking. Kymographs of the rings labeled with Rlc1-tdTomato were generated to determine both the start and end of ring closure. The customized plug-in Ring Intensities Measurement-2015 was used to measure the GFP-LA fluorescence during the constriction of rings that went on to complete constriction (Courtemanche et al., 2016). The background fluorescence was calculated separately for each cell used for GFP-LA measurement. The corrected GFP-LA fluorescence was then converted to number of actin molecules based on the calibration as described (Morris et al., 2019).

The number of actin filaments in a cross-section of the contractile ring was calculated by dividing total length of actin filaments with the circumference of a cell. Total length of actin filaments (µm) is calculated as total number of actin molecules divided by 370. The circumference of wild-type and *adf1-M3* cells was estimated at 10 and 13 µm respectively (Chen and Pollard, 2011). The center-to-center spacing between actin filaments is assumed to be 15 nm (Kanbe et al., 1989; Swulius et al., 2018) in estimating the width of a contractile ring.

### Western Blots

For probing the expression level of GFP-Lifeact, 50 ml of exponentially growing cells were collected by centrifuging at 3000g for 5 min. The pellet was resuspended in 300 µL of lysis buffer containing Halt™ protease inhibitor cocktail (#1862209, Thermo-Fisher Scientific) and frozen at-20 °C until lysis. The cells were lysed using BeadBug microtube homogenizer (Benchmark Scientific). Proteins were resolved using Mini-PROTEAN TGX™ Precast SDS-PAGE gels (#4561084, BioRad). The gels were either stained with Coomassie blue or transferred to PVDF membrane (Millipore) for immunoblots. Membranes were blotted with 1:1000 dilution of primary anti-GFP (#11814460001, Sigma-Roche) over night, followed by 1:5000 dilution of HRP-linked secondary antibody (#sc-516102, Santa Cruz) for 2 h. The blot was developed using SuperSignal™ West Pico PLUS chemiluminescent substrate (#34577, Thermo-Fisher Scientific).

## Acknowledgements

Research reported in this publication was supported by National Institute of General Medical Sciences of the National Institutes of Health under award numbers R01GM026132 to TDP and R15GM134496 to QC. The content is solely the responsibility of the authors and does not necessarily represent the official views of the National Institutes of Health. The authors thank Debatrayee Sinha and Abhishek Poddar for their technical assistance.

## Figures and Figure legends

**Supplemental Figure S1:**
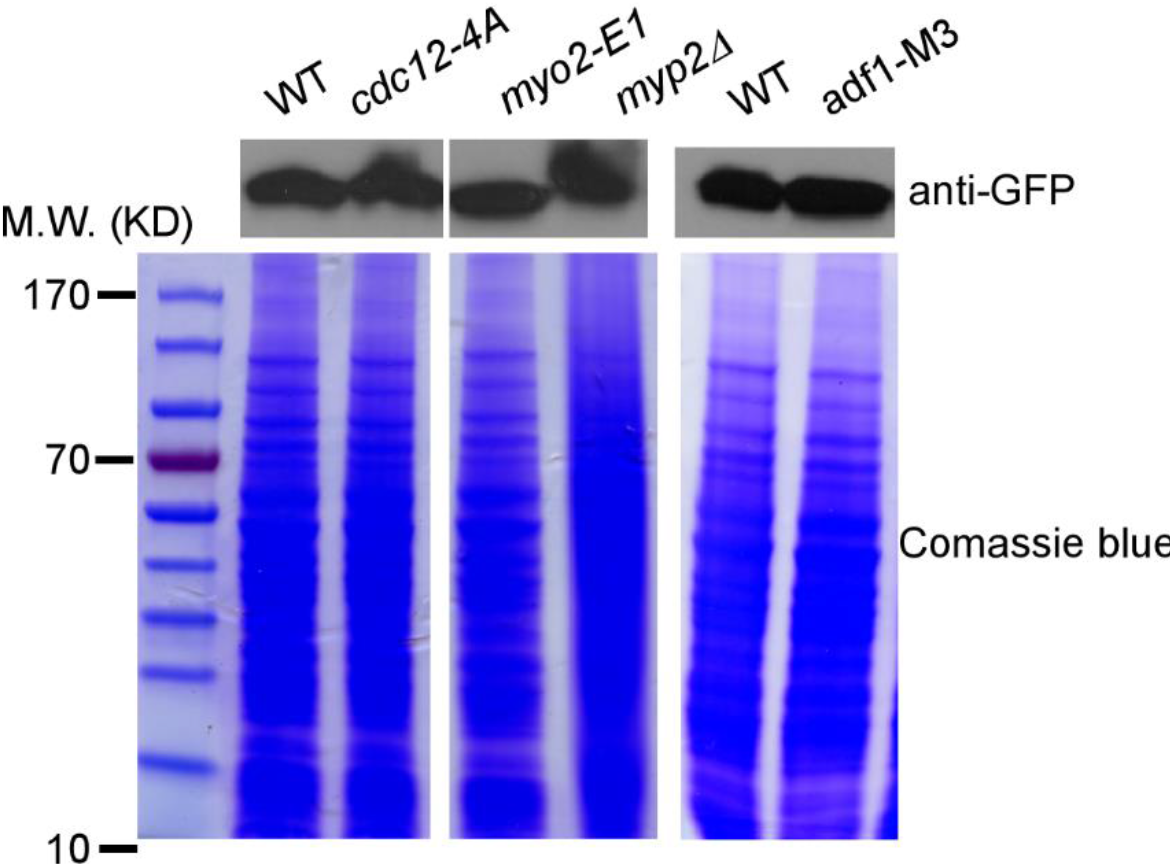
Expression of GFP-Lifeact. Top: Anti-GFP blot of the whole cell lysate from wild-type, formin mutant and myosin mutants (left) and wild-type and cofilin mutant cells (right). Bottom: Coomassie blue-stained SDS-PAGE gel of the whole cell lysate as loading control.

**Supplemental Figure S2:**
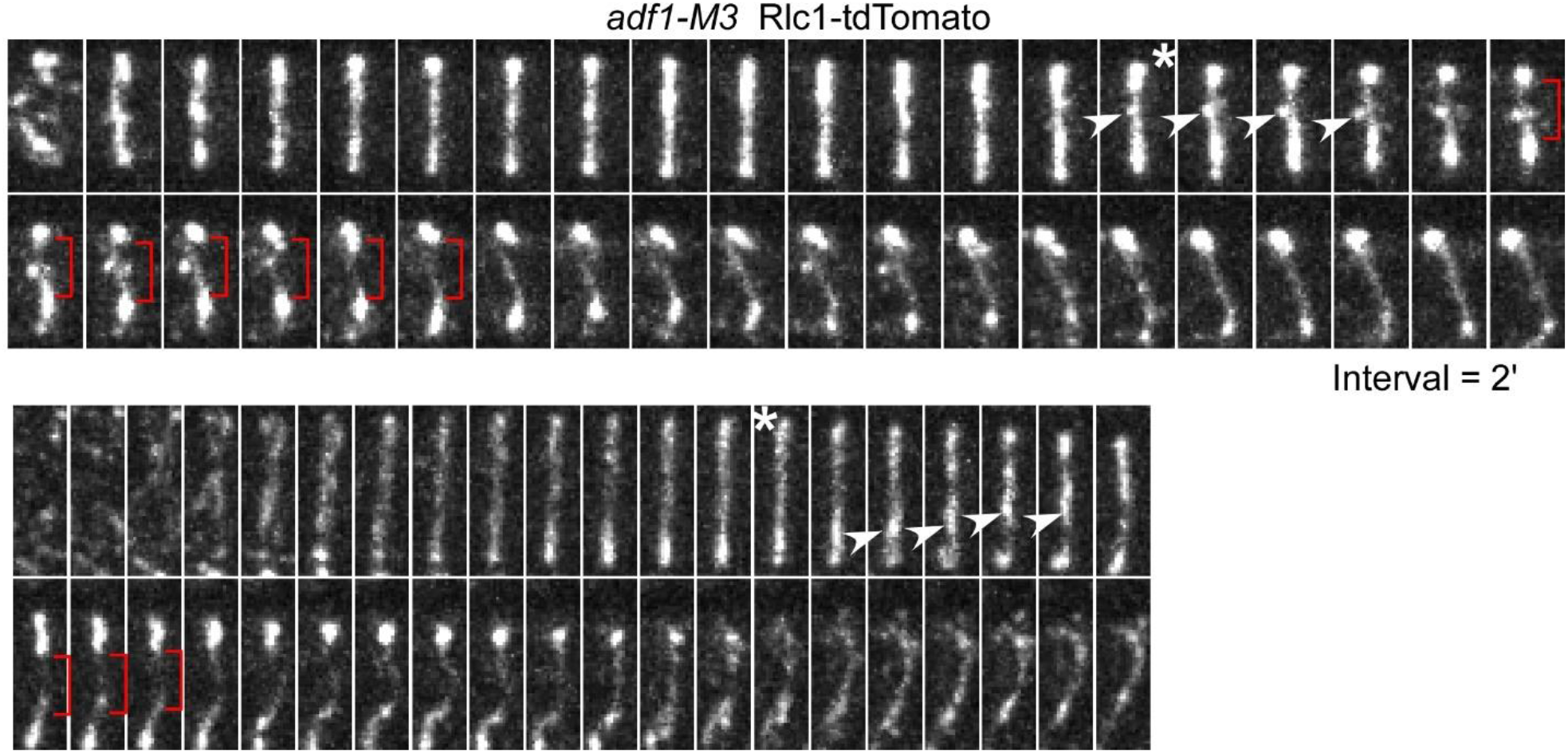
The contractile rings that failed to constrict in the cofilin mutant cells. Time-series micrographs of two *adf1-M3* cells expressing Rlc1-tdTomato. Asterisk: start of the contractile ring constriction. Arrowhead: fragmentation of the ring. Bracket: enlarging gap in the contractile ring. Such fragmenting rings represent 30% of the contractile rings among the cofilin mutant.

